# Long-term Tolerance to Islet Transplantation via Targeted Reduction of beta cell-specific T cells

**DOI:** 10.1101/2024.09.02.610863

**Authors:** Michael Kotliar, Eileen E. Cianciolo, Duc Hung Pham, Kaitlin R. Carroll, Artem Barski, Michael B. Jordan, Jonathan D. Katz

## Abstract

Type 1 diabetes (T1D) results from insulin insufficiency due to the loss or dysfunction of pancreatic beta cells following T cell-mediated autoimmune attack. Currently the only long-term therapy is daily exogenous insulin replacement. The ideal curative approach is the durable restoration of functional islets via transplantation. To date the limiting factors impeding realization of this goal is the lack of a cost effective and limitless source of high-quality islets suitable for transplantation and the ability to provide long-term islet graft acceptance without prolonged need for deleterious immunosuppression. Ongoing clinical trials are testing islets derived from human induced pluripotent stem cells (iPSC); however, long-term acceptance of islet graft will require a effective therapeutic strategy to prevent engrafted islet destruction by pre-existing islet-antigen specific T cells. Here we demonstrate in the NOD mouse model for T1D that autologous islet graft acceptance can be achieved by the targeted elimination of (re)-activated islet-reactive CD4^+^ and CD8^+^ T effector (Teff) cells in the initial post-transplantation period by using a short-acting, combination therapy that results in the elimination of islet-reactive Teff cells by exacerbation of their natural DNA damage response (DDR) to drive apoptosis while at the same time maintaining endogenous Treg cells.

**Article Highlights:**

- Activated beta-cell reactive CD4^+^ and CD8^+^ T effector cells undergo a profound DNA-damage response which is targetable by small molecule inhibitors of the p53 and cell cycle pathways that lead to apoptosis.
- The use of a combination of MDM2 and WEE1 inhibitors, which termed “p53 potentiation with checkpoint abrogation” (PPCA), conferred significant therapeutic efficacy in treating mouse models of new onset T1D.
- Specific targeting of these T effector cells by PPCA results in a loss of inflammatory T cell subsets, notably proliferation CD4^+^ Th0 and Th1 subsets and CD8^+^ T effector memory cells, as determined by single cell RNA-seq studies with the preservation of T regulatory cells.
- When autologous islet grafts are given to established diabetic NOD mice, a single course of PPCA results in long-term islet graft acceptance, restoration of normoglycemia and loss of beta cell specific CD4^+^ and CD8^+^ T cells.
- PPCA shows promise as a potential means of estimating islet graft tolerance in T1D recipients of islet graft transplantation.

## Introduction

Type 1 diabetes (T1D) is a T cell-driven autoimmune disease that arises in genetically predisposed individuals influenced by environmental and stochastic events (1), resulting in the dysfunction and death of insulin-secreting β-cells of the pancreas. The challenge in preventing or curing T1D lies in the clinically silent nature of its pathogenesis, with symptoms only appearing after a substantial proportion of a patient’s β-cells cease insulin production or are outright destroyed (1). For a century now, exogenous insulin replacement remains the only life-sustaining therapy, and despite substantial advances in insulin production, formulation, and delivery, as well as improvements in glucose monitoring, only a third of T1D patients achieve sufficient glucose control to avoid secondary sequelae such as peripheral neuropathy, micro- and macro-vasculopathies, retinopathy, and renal failure (2).

T1D is driven primarily by autoreactive CD4^+^ and CD8^+^ T cells (1; 3). CD8^+^ T cells damage β-cells directly to induce apoptosis via perforin and granzyme release and interferon-γ (IFN-γ) secretion (3; 4). T helper (Th)-1 and Th17 CD4^+^ T cells (and their plastic intermediaries) mediate β-cell death by secreting IFN-γ, tumor necrosis factor-α (TNF-α), and interleukin-17 (IL-17), which are cytocidal to β-cells and induce and activate M1 macrophages to secrete additional proinflammatory cytokines in a feedforward loop (4; 5).

Regulatory CD4^+^ T cells (Tregs) are critical antagonists of effector T cells (Teff) driven T1D pathogenesis. Studies in T1D patients suggest when compared to healthy controls their Tregs lack suppressive function (6) or produce IFN-γ (7), or that their Teff cells exhibit intrinsic resistance to Treg suppression (7–9). Nonetheless, the preservation or enhancement of Treg cell function in the context of T1D is clearly of clinical benefit (10; 11).

β-cells isolated from patients with new onset T1D regained ex vivo glucose-mediate insulin production when removed from their diabetogenic environments (12), suggesting that in the absence of stressors β-cells might regain functionality *in vivo*.

Islet replacement is the therapeutic strategy with the greatest likelihood of providing a durable long-term cure, as it would not only provide a new source of insulin production but also likely relieve metabolic stress on any residual endogenous pancreatic β cells. Islet transplantation and replacement continues to progress (13–15); however, the central problems of islet replacement remain the availability of islets and the long-term survival and function of the graft facing auto- and allo-immunity (13). The feasibility of producing and transplanting functional β-cells – especially from iPSC – shows promise, keeping them functional in an immunologically hostile environment surrounded by pre-existing insulin-reactive lymphocytes remains an ongoing struggle (16; 17). To date, most efforts have focused on generalized immune suppression, complete ablation of immune compartments, or enhanced immune regulation (18).

Given that T1D arises in part due to a failure in peripheral tolerance, it seems logical that depleting the pre-existing pathogenic T cells is the most direct and effective way to halt disease progression (19). Studies in NOD mice suggest that both decreasing the frequency of autoreactive Teff cells or increasing the frequency and function of Tregs each provide protection from T1D, reducing autoreactive Teff cells shows greater efficacy (20–23). The challenge, however, is how to deplete only islet-reactive cells without impacting protective immunity.

Recently, we demonstrated that one potential powerful means of eliminating islet-reactive Teff cells is via the manipulation of the natural DNA damage response (DDR) pathway intrinsic to CD4^+^ and CD8^+^ Teff at the time of diabetes onset (24). Activated Teff cells undergo high metabolic and proliferative stress and exhibit a profound DDR (25), with the phosphorylation of DNA damage sensory proteins, ataxia telangiectasia mutated (ATM) and ATM and RAD3 related (ATR), the activation of their downstream targets CHK1 and CHK2 and their upregulation of the kinase WEE1, all of which inhibit CDK1 and CDK2 and cell cycle progression (26). ATM and ATR also act by phosphorylating and activating p53, a transcription factor, which in turn causes the transcription of p21 that inhibits CDK2 and causes G1 arrest (27). Ordinarily, p53 is kept during homeostasis at relatively low levels by its negative regulator MDM2, an E3-ubiquitin ligase (26; 28). When p53 is phosphorylated, it dissociates from MDM2 and accumulates in the cell (26), where its transcriptional activity differs depending on its intracellular levels; at lower levels it mediates G1 arrest via p21 transcription, but at higher levels it induces the transcription of pro-apoptotic molecules such as BAX, PUMA, and NOXA (26; 28). Thus, p53 sits at the nexus between damage repair and apoptosis.

The intrinsic DDR in activated Teff cells, therefore, puts them at the brink of apoptosis and makes them exquisitely susceptible to minor perturbations in activated p53 levels and downstream regulators of cell cycle control (25). We demonstrated that exposing activated Teff cells to short-acting, small molecule inhibitors that target CHK1/2 or WEE1 interrupts cell cycle regulation and DNA damage repair and when coupled with a small molecule that stabilizes phospho-p53, forces p53-mediated apoptosis (25) thereby eliminating activated Teff cells. We showed that this combination, which we have termed “p53 potentiation with checkpoint abrogation” (PPCA), conferred significant therapeutic efficacy in treating mouse models of not only new onset T1D (24; 28) but also other diseases caused by aberrant T cell activation, multiple sclerosis (experimental autoimmune encephalomyelitis; EAE) and hemophagocytic lymphohistiocytosis (25; 29).

Herein we demonstrate at a single-cell level that PPCA preferentially targets the loss of proliferating intra-islet early CD4^+^ Teff cells (Th0), Th1 CD4^+^ cells, and CD8^+^ T effector memory (Tem) cells. Moreover, the Th1 and Th17 cells that persist after PPCA therapy have reduced inflammatory cytokine production, while the colocalized Treg cells retain both their numeric and functional ability to regulate T1D. Critically, a single round of PPCA treatment in the initial window post-islet transplant results in significant long-term autologous islet graft survival and restoration of euglycemia diabetic NOD graft recipients suggesting that focused PPCA therapy at the time of synchronized reactivation of pre-existing β-cell-specific CD4^+^ and CD8^+^ Teff cells is highly efficacious and results in long-term islet graft survival.

## Research Design and Methods

### Mice

NOD/ShiLtJ (NOD), NOD.129S7(B6)-Rag1^tm1Mom^/J (NOD.Rag1^-/-^), NOD.129X1(Cg)-Foxp3^tm2Tch^/DvsJ (NOD.Foxp3^eGFP^) were purchased from the Jackson Laboratory, bred, and maintained under specific pathogen-free conditions in accordance with institutional animal care guidelines at Cincinnati Children’s Medical Center Vivarium.

### Blood Glucose Assessment

Prediabetic NOD mice were monitored beginning at 10 weeks of age for clinical signs of prediabetes (hyperglycemia, assessed by non-fasting blood glucose testing). Diabetes onset was determined via two consecutive blood glucose readings of ≥200 mg/dL. Mice were monitored tri-weekly from this point until they developed end-stage disease (blood glucose≥600 mg/dL), lost more than 20% body weight, established long term normoglycemia for at least 60 days, or died.

### Drug Treatments

All chemotherapeutics were administered intraperitoneally and were dosed as follows: AZD1775 (WEE1i; Chemietek), 40 mg/kg; AZD7762 (CHKi; Selleck Chem), 25 mg/kg; nutlin-3 (MDM2i; Cayman Chemical), 50 mg/kg. The vehicle was made as previously described (24).

### MHC Tetramer Staining and Flow Cytometry

Single-cell suspensions of spleen and pancreatic lymph nodes were separated by density gradient centrifugation using Lympholyte-M and incubated with tetramer (NIH Tetramer Core). Cells were then stained for surface expression of the indicated antibodies. For intracellular staining, cells were fixed with the FoxP3 fixation/permeabilization buffer and permeabilized with permeabilization buffer. All antibodies were purchased from BioLegends. Flow data was collected using an Fortessa system using FACS Diva software or Cytek Aurora flow cytometer and analyzed using FlowJo™ Software for Windows Version 10.10.0. Ashland, OR: Becton, Dickinson and Company; 2023.

### Isolation of Pancreatic Lymphocytes

Single-cell suspension of pancreatic cells was prepared using the Miltenyi gentleMACS Dissociator, filtered through a 70um cell strainer, and prepared according to the Miltenyi Debris Removal protocol. Isolated lymphocytes were then stained in accordance with the methods listed above.

### Histological Scoring/Immunofluorescence

Islet kidney grafts were formalin fixed and paraffin embedded. Sections were stained with antibodies to insulin and glucagon (Abcam) followed by the indicated secondary antibody (Abcam). Imaged using the CCHMC Bio-Imaging and Analysis Facility. Images were analyzed using NIS Elements Imaging Software.

### T Cell Isolation and Enrichment

T cells were prepared for adoptive transfer from NOD.Foxp3^eGFP^ mice and isolated as described above, followed by sort-purification on a Sony MA900 flow sorter. Treg cells were identified using eGFP and CD4. Teff cells were sorted based on CD4 and CD44 expression and the lack of eGFP. 10^7^ Teff cells and indicated numbers of purified Treg cells were injected via retro-orbital intravenous injection in 250 µl DMEM into NOD.Rag1^-/-^ mice. To assess intracellular cytokine production mice may be treated with brefeldin A (100 μg) 6 hours before harvest.

### Next-Generation scRNA-Seq and Analysis

Single-cell RNA sequencing (scRNA-Seq) was conducted using Chromium Single Cell 3’ Library & Gel Bead Kit v3 and run on a 10X Genomics Chromium platform. PolyA mRNA purification, quality control, and next-generation sequencing of 8 samples (1 active T1D, 1 onset T1D, 3 from PPCA, and 3 from vehicle treatment group) were performed by the University of Cincinnati Genomics, Epigenomics and Sequencing Core (GESC). To quantify gene expression at a single-cell level, FASTQ files obtained from GESC for each sample were individually aligned to pre-built mouse reference genome indices (GENCODE vM23 / Ensembl 98; https://www.10xgenomics.com/support/software/cell-ranger/latest/analysis/inputs/cr-3p-references using the 10x Genomics Cell Ranger 6.1.2 (30) command-line tool. The resulting feature-barcode matrices (one per sample), each containing Unique Molecular Identifier (UMI) counts per gene and cell combination, were merged with a custom R script for joint analysis in subsequent steps. Cell barcodes were updated to reflect their samples of origin. All single-cell data analysis steps described below were performed in SciDAP (https://SciDAP.com) following the instructions outlined by (31) (https://doi.org/10.1101/2024.02.28.582604).

“Single-Cell RNA-Seq Filtering Analysis” pipeline was used for removing low-quality cells. The following Quality Control (QC) filtering thresholds were applied to all samples: minimum of 1000 reads per cell, maximum of 5% of reads mapped to mitochondrial genes, Additionally, the thresholds for the minimum and maximum number of genes per cell were selected independently for each sample as follows: 1,085 – 3,000 for active T1D, 890 – 2,575 for onset T1D, 960 – 2,800 for vehicle treatment on day 2, 850 – 2,450 for PPCA treatment on day 2, 820 – 2,600 for vehicle treatment on day 3, 760 – 2,550 for PPCA treatment on day 3, 840 – 2,500 for vehicle treatment on day 4, and 650 – 2.170 for PPCA treatment on day 4.

Dimensionality reduction and samples integration were conducted using “Single-Cell RNA-Seq Dimensionality Reduction Analysis” pipeline. The workflow was run with 40 dimensions and 3,000 highly variable genes. To match shared cell types across multiple samples, as well as to correct for technical differences, the integration method was set to “seurat”. Thus, the cross-dataset pairs of cells that were in a matched biological state were used as integration anchors (32). The percentage of reads mapped to mitochondrial genes, as well as the expression levels of Trbc1 and Trbc2 genes, were regressed out as the confounding sources of variation.

In the next step “Single-Cell RNA-Seq Cluster Analysis” pipeline was executed to group cells by gene expression similarities. Target dimensionality and clustering resolution parameters were set to 30 and 0.3, respectively. Using the known hallmark genes, the following cell types were identified and excluded from the subsequent data analysis steps: NK cells (Klra1, Ncr1), dendritic cells (Myb, Gpr183), exocrine cells (Ctrb1, Prss2, Cela2a), monocytes (Lgals3, Tmem176a, Tmem176b), and B cells (Cd19, Cr2, Ms4a1). Additionally, a cluster with both Cd4^+^ and Cd8a^+^ cells and without any other distinct gene markers was removed as doublets.

After removing cell types of not particular interest, the “Single-Cell RNA-Seq Dimensionality Reduction Analysis” and “Single-Cell RNA-Seq Cluster Analysis” pipelines were rerun with 30 and 15 dimensions, respectively. Clustering resolution was set to 0.22 to achieve optimal separation between the expected cell types. Cluster annotation was made on the basis of the following marker genes: CD4 Naïve (Cd4, Ifngr2, Foxp1), CD8 Tcm (Cd8a, Ccr7, Sell), CD8 Trm Tem (Cd8a, Ccl5, Itga1), CD4 Th1 (Cd4, Ifi27l2a, Ifit1, Ifit3, Igtp, Iigp1, Stat1), CD4 Treg (Cd4, Foxp3, Ctla4), CD4 Th17 (Cd4, Tbx21, Rora, Rorc, Ifng), and CD4 Th0 (Cd4, Cd69, Nfkbid, Nfkb1).

Gene set enrichment analysis was performed using GSEAPreranked tool from the GSEA 4.0.3 (33) desktop application run with default parameters. Hallmark gene sets v7.5 from the Human Molecular Signatures Database (MSigDB) (34) were selected as the reference, whereas the ranked gene lists were obtained from the “Single-Cell RNA-Seq Differential Expression Analysis” pipeline run for each cell type and timepoint (excluding day 1) between the PPCA and vehicle treated mice. Mouse genes were converted to human equivalents using biomaRt R package (35). P-values and log-fold changes were calculated by FindMarkers function with default parameters. The rank of each differentially expressed gene was assigned based on its log-transformed p-value and the direction of change (positive for upregulated and negative for downregulated genes). Datasets have been deposited with the following accession number: GSE262644 at: https://www.ncbi.nlm.nih.gov/geo/query/acc.cgi?acc=GSE262644

### Islet Transplantation and nephrectomy

Diabetic mice were treated for 21 days prior to surgery with 3-4 LinBit implants (LinShin Canada, Inc.) based on body weight to provide sustained insulin delivery and normoglycemia. Islet isolations and transplants were performed as previously described (24; 36); briefly, recipient NOD mice will be anesthetized with isoflurane (2-3% then too effect), the site is prepared using hair removal and proper antiseptic technique. A small incision will be made on the left flank, just large enough to expose the left kidney. A small incision will be made to the kidney capsule, and the harvested islets will be injected underneath the kidney capsule through a small tube. The incision to the peritoneal membrane will be closed using absorbable sutures, and the incision to the skin will be closed using surgical staples (which either fall out naturally or are removed after 1 week). Post surgical analgesics are given and post op pain monitored. NOD mice received ∼300 islets isolated from NOD.Rag1^-/-^ mice as a source of autologous islets without adaptive immune infiltrate. For mice that underwent post-transplant nephrectomy, the left flank incision was reopened, the left kidney was exposure and venal and atrial blood flow and ureter cauterized and the kidney was freed from the abdomen. The incision to the peritoneal membrane was closed using absorbable sutures, and the incision to the skin was closed using surgical staples. After nephrectomies mice developed hyperglycemia with 24h indicating that the engrafted kidney was responsible for maintaining euglycemic control.

### Statistical Analysis

All statistical analyses were performed using one- or two-way ANOVA, log-rank Mantel-Cox, or multiple t-test using GraphPad Prism version 10.2.0 for Windows.

## Results

Previously, we established that combining an inhibitor of MDM2 such as Nutlin-3 or idasanutlin (here after MDM2i) with an inhibitor of cell cycle such as AZD1775, which targets WEE1 (here after WEEi) or AZD7762, which targets CHK1 and 2 (here after CHKi), effectively drove apoptotic elimination of both activated CD4^+^ and CD8^+^ Teff in a therapeutic modality we termed PPCA (24; 25). Importantly, this treatment acts on rapidly proliferating Teff cells. Naïve T cells, in G0, with little or no DNA damage (25) and Tregs, which favored primarily aerobic glycolysis, and utilize catabolic fatty acid oxidation (FAO) and oxidative phosphorylation with minimal DDR (37; 38), are largely resistant to PPCA. Quiescent memory T cells with their slower homeostatic turnover rates and use of FAO and oxidative phosphorylation are like Tregs not targets of PPCA therapy. When PPCA therapy is applied to NOD mice at the onset of T1D they significant autoreactive Teff cells are depleted, halting further damage to β-cells, sustaining insulin production, and allowing the mice to remain normoglycemic without exogenous insulin administration for significantly longer than vehicle-treated controls (24). Additionally, PPCA significantly shifts the balance of β-cell-specific Teff and Tregs cells; decreasing islet-specific CD4^+^ and CD8^+^ Teff cells while preserving Tregs cells.

To determine to what extent PPCA favored the elimination of specific T cell subsets, especially those known to drive T1D, namely CD4^+^ Th1 and Th17 cells and CD8^+^ Tem cells, as well as determine their loss rate and phenotypic change under PPCA therapy, we performed a longitudinal study of the effects of PPCA on pancreatic infiltrating T cells at the onset of T1D in NOD mice. Wild type female NOD mice were aged and allowed to develop spontaneous new onset T1D (200-250 mg/dl, blood glucose) at which time, day 1, they were randomly assigned to either the PPCA or vehicle treatment group, whereupon the mice were treated with PPCA (WEE1i + MDM2i) or vehicle daily for three days (day 2-4). Each day, prior to the next round of treatment randomly selected mice from each group were sacrificed, and their pancreata harvested and the CD45^+^ CD3^+^ T cells were isolated first by gradient centrifugation and then by flow cytometric cell sorting. The resulting T cells were then subjected to single-cell RNA sequencing (scRNA-Seq) allowing us to exam the dynamic changes in T cell subsets at a single-cell level over the course of our 3-day PPCA treatment; new onset (day 1) through one (day 2), two (day 3) or three (day 4) rounds of PPCA therapy. As a control, we additionally interrogated T cells from the NOD mouse that have already developed active T1D. Adding these data to the analysis increased the accuracy of dataset integration and cell type assignment. Gene expression profiles from all individual samples were estimated at a single-cell level and merged into a single feature-barcode matrix for joint analysis. We identified seven distinct T cell subsets within the pancreas of NOD mice at the time or onset and treatment, **Figure 1A**, that included CD8^+^ T effector memory (CD8^+^ Tem) and T central memory (CD8^+^ Tcm) subsets and CD4^+^ naïve T cells (CD4^+^ naïve), early proliferating Th0 effectors, CD4^+^ Th1, CD4^+^ Th17, and CD4^+^ Treg cell subsets. Of note, we did not find any evidence for CD4^+^ Th2 cells consistent with previous data suggested that Th2 cells play little to no role in T1D development (4; 39). We then performed randomized down sampling on individual dataset to normalize each to 5,757 cells (the size of the smallest dataset) so that the dataset could be equally compared to each other to assess quantitative and qualitative changes during PPCA treatment. As depicted in **Figure 1B** and quantified in **Figure 1C**, over the three day course of PPCA, the numbers of CD4^+^ Th0, and Th1 cells significantly declined, as did the CD8^+^ Tem subset, when compared to the vehicle treated mice. Importantly, the numbers of Treg cells were largely unchanged. These data suggest that PPCA treatment leads to progressive loss of key inflammatory effector populations.

**Figure 1:**
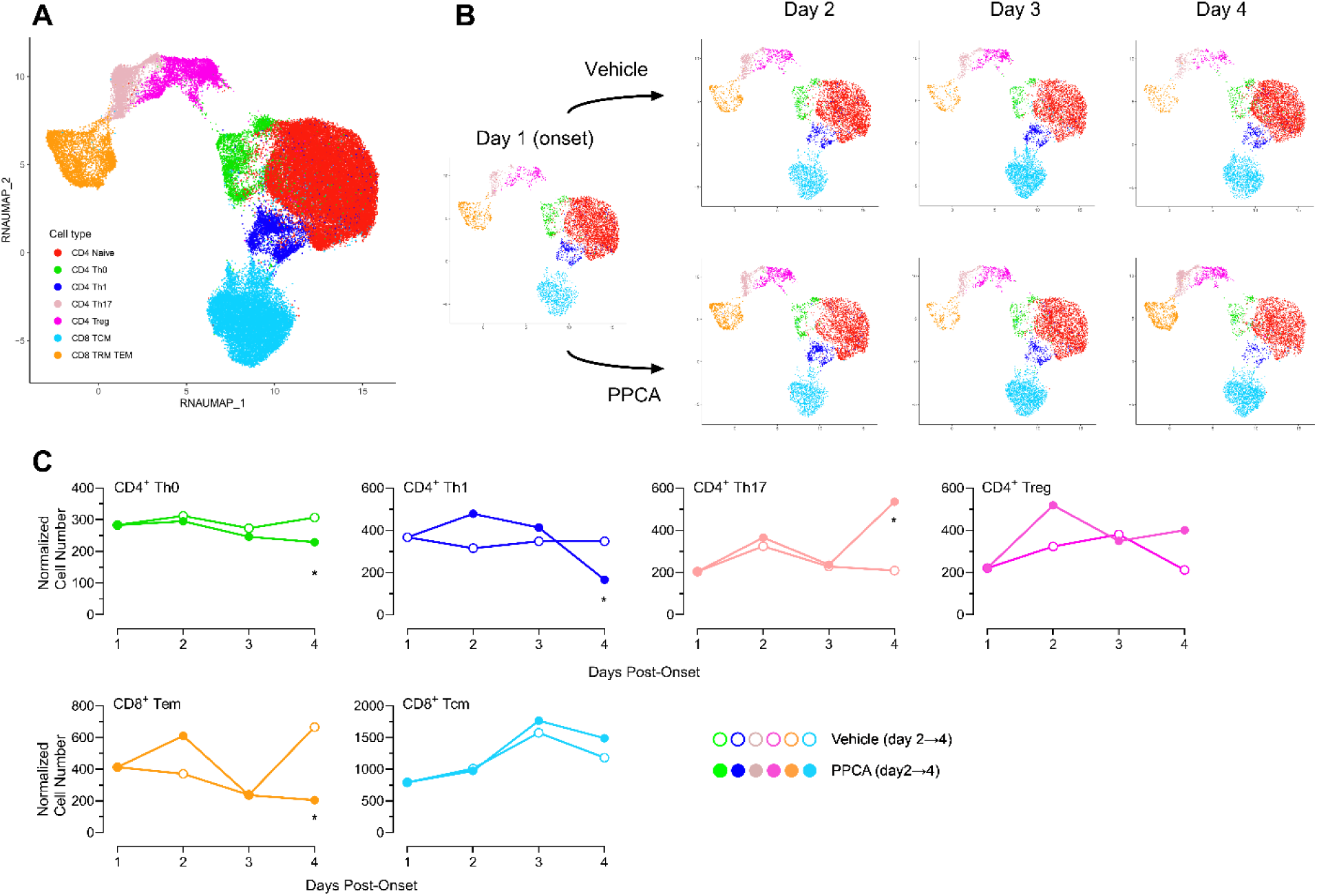
PPCA therapy depletes proliferating CD4^+^ Th0, CD4^+^ Th1 and CD8^+^ Tem cells while sparing CD4^+^ Treg cells as revealed by scRNA-Seq. Diabetic NOD female mice, n=9, were treated with PPCA or vehicle for 3 consecutive days post new onset T1D and pancreatic lymphocytes were isolated in a subset of each group 6 hours after each round of therapy. **A.** Clustered and annotated UMAP plot of integrated with Seurat samples from onset T1D, PPCA and vehicle treatment groups. 7 distinct cell types were identified on the basis of the known hallmark genes. **B.** Clustered and annotated UMAP plots split by dataset. Cell counts are down sampled to the size of the smallest dataset saving the proportion of the cells within each cell type and allowing for direct comparison between the samples. **C.** Normalized to the size of the smallest dataset cell counts per sample for each of the identified cell types enumerated and compared for dynamic modulation over time. Differences in normalized cell counts were compared using multiple linear regression analysis (Shapiro-Wilk) using GraphPad Prism (Version 10.2.3). * Denotes p≤0.05.

PPCA treatment not only reduced the overall numbers of these critical inflammatory T cell subsets, but also dynamically altered the gene expression profiles of the remaining cells within each subset over time under treatment pressure. Using GSEA analysis to compare the gene expression dynamics within each subset between the PPCA and vehicle treated mice, we identified distinct global and subset-specific dynamic changes. For the first two days of PPCA therapy, we observed in all subsets a positive, but diminishing, global enrichment score indicative of a response to ongoing type 1 and type 2 interferon driven inflammation, **Figure 2**. In the presence of PPCA, the Th0 and Th1 subsets showed a statistically significant (p≤0.01) shift away from an inflammatory gene expression profile and toward one associated with p53, cell stress and apoptosis. While the shift in the Th17 population was less dynamic, by Day 4, they too showed enhanced response to p53 (**Figure 2**). The Treg subset under PPCA therapy surprisingly exhibited a decrease in IL-2 signaling initially but then rebounded by Day 4. This paralleled to some extent the gene expression dynamics of the Th0 population leaving open the possibility that some of the change in the Th0 subset on Day 3 and 4 may represent the induction of new Treg cells, which is consistent with maintenance of a steady state of Tregs over the treatment period while Th0 cells were lost, **Figure 1**.

**Figure 2:**
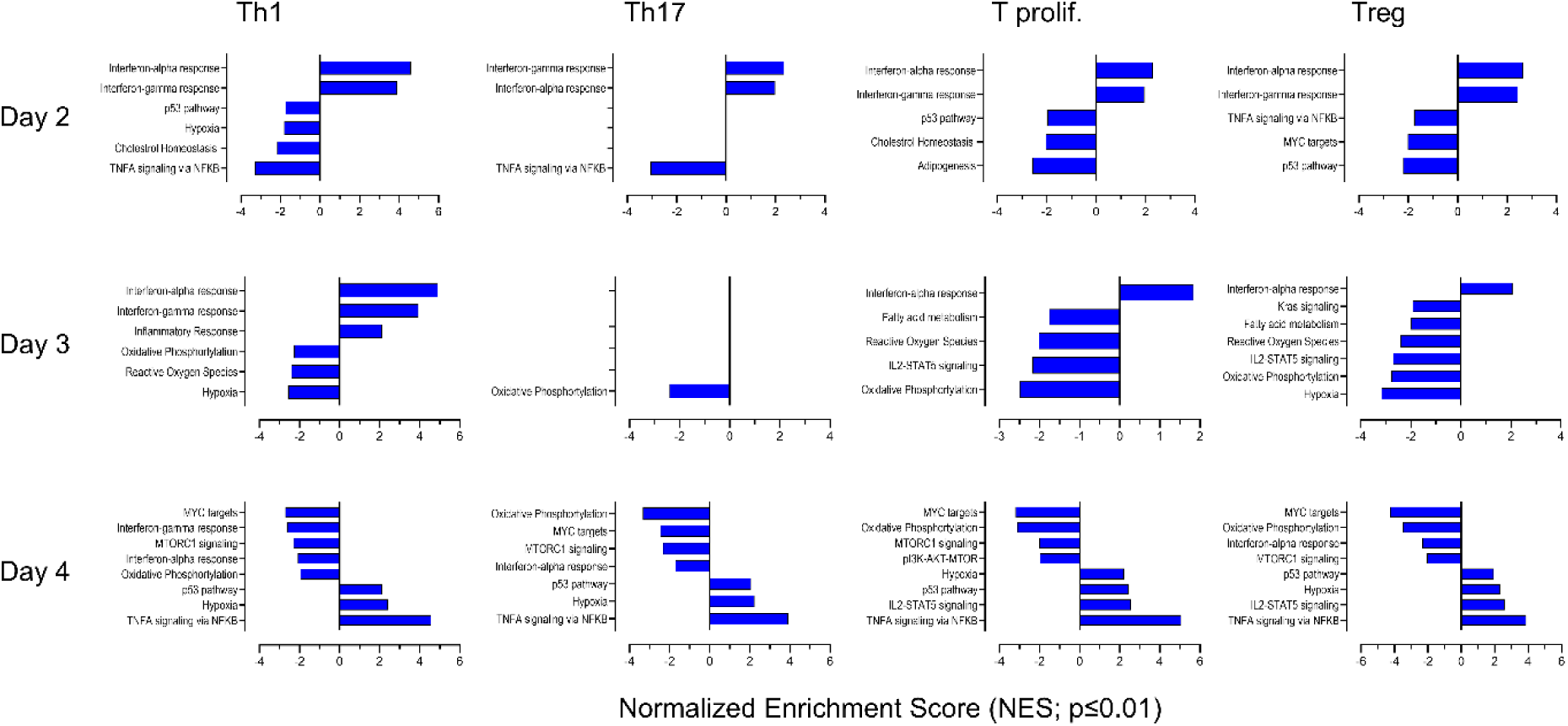
In the presence of PPCA, the Th0 and Th1 subsets showed statistically significant shift away from an inflammatory gene expression profile and toward one associated with p53, cell stress and apoptosis. For cell clusters in each dataset gene set enrichment analysis was performed comparing the PPCA treated verses the vehicle treated mice at the same timepoint and compared to curated rank ordered gene list comparing genes positively correlated with PPCA treatment (>0) versus those negatively correlated (<0) on Day 2 through Day 4); bar colors assigned based on correlation of Th1 cells on Day 2. Normalized Enrichment Scores (NES) from those lists with a p≤0.01 are shown. As PPCA treatment progressed, Th1 and Th0 cells showed a reversal of correlation from proinflammatory type 1 and type 2 interferon response towards one associated with p53 signaling, cell stress (hypoxia) and altered cell metabolism and apoptosis. Th17 were less dynamic at Day 2 and 3 but paralleled the Th1 cells on Day 4. Treg cells showed enhanced IL-2 responsiveness on Day 4 even as proinflammatory responses waned.

While PPCA therapy clearly reduced the numbers of proinflammatory T cell subsets in the pancreas, Th1 and Th17 do remain after therapy yet, these mice had prolonged T1D remission and delayed or blunted transition to end stage T1D (**Figure 1** and (24)). While an inversion of the Teff to Treg balance can explain the enhanced control of these residual Teff cells, an alternative, but not mutually exclusive, explanation exists that PPCA therapy reprograms or favors the survival of less pathogenic Teff cells. Intriguingly, we found that intrapancreatic Th1 cells from PPCA but not vehicle treated mice upregulated Usp18 (40; 41), the ISG56/IFIT1 gene family (42) (Ifit1 and Ifi3b), as well as Rsad2 (43), and Uba52 (44), **Figure 3A**, all of which are induced by type 1 interferon signaling in concert with endogenous DNA damage but exert opposing regulatory effects on inflammation and subsequent interferon gene programs, including reduced cytokine production. Notably, Usp18 acts as a negative regulator of IFN-γ and IL-17 production, as loss of Usp18 was associated with augmented IFN-γ production and enhanced Th17 maintenance and IL-17a and IL-17f production (45; 46). To assess if the upregulation observed in PPCA-treated Th1 cells resulted in altered levels of IFN-γ production, we treated new onset T1D NOD mice with PPCA or vehicle control for three days, and with brefeldin A for last 6 hours prior to harvesting pancreatic lymph node cells to directly assessing *ex vivo* intracellular IFN-γ production by flow cytometry. As shown in **Figure 3B** and **3C**, CD4^+^ CD44^+^ T cells from the draining pancreatic lymph nodes of PPCA-treated mice had diminished IFN-γ production compared to vehicle controls, consistent with not only a reduced number of Th1 cells, but also a significant reduction in localized IFN-γ producing cells after PPCA intervention.

**Figure 3:**
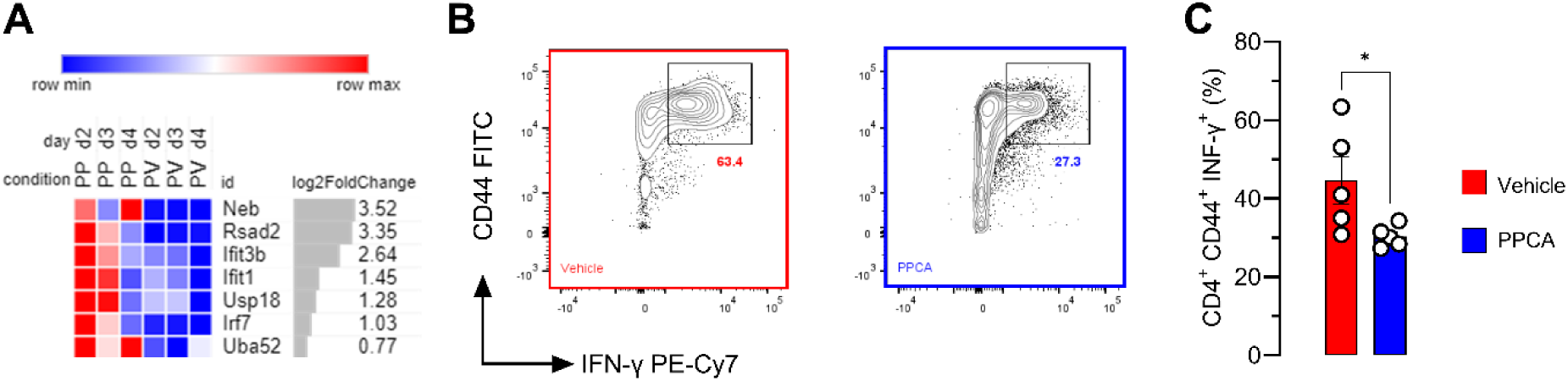
Upregulation of type 1 interferon induced counter regulatory genes that down modulate IFN-γ expression in residual Th1 cells following PPCA therapy. **A.** Treatment of Th1 cells with PPCA for 3 days results in upregulation of Type 1 interferon driven genes associated with counter-regulation of inflammatory responses. Analysis of aggregated pseudo-bulk of the logFC direction of PP (PPCA treated pancreatic cells) versus PV (vehicle treated pancreatic cells). Log2 Fold Change on right. **B.** Representative flow cytometric analysis of intracellular IFN-γ production from PPCA and vehicle control treated NOD mice shows diminished IFN-γ production in PPCA treated CD4^+^ CD44^+^ Teff cells. **C.** Cumulative expression of IFN-γ in CD4^+^ CD44^+^ Teff cells. * Denotes p<0.05, two-tailed paired t test.

There was no significant loss of Treg cells upon PPCA treatment here or in previous studies (24), however, to directly assess if PPCA therapy altered the suppressive functionality of Treg cells, we treated new onset T1D NOD.Foxp3-eGFP mice with PPCA or vehicle control for 3 days and then harvested and sort-purified (GFP^+^) Treg cells, as detailed in **Sup Figure 1**, from each group and co-transferred defined numbers of these Tregs cells along with a fixed dose of 1 x 10^7^ Treg-depleted (GFP^-^) Teff cells (from vehicle treated T1D NOD.Fox3-eGFP mice) into NOD.Rag1^-/-^ recipients. As expected, we found no functional variation, at any Teff to Treg ratio, in the ability of PPCA-treated Treg cells to regulate the T1D induced in NOD.Rag1^-/-^ recipients by NOD Teff cells, **Figure 4**. At all three Treg cell ratios, we observed no significant ratio-dependent reduction in T1D regulation affected by Treg cells from either PPCA or vehicle control treated mice.

**Figure 4:**
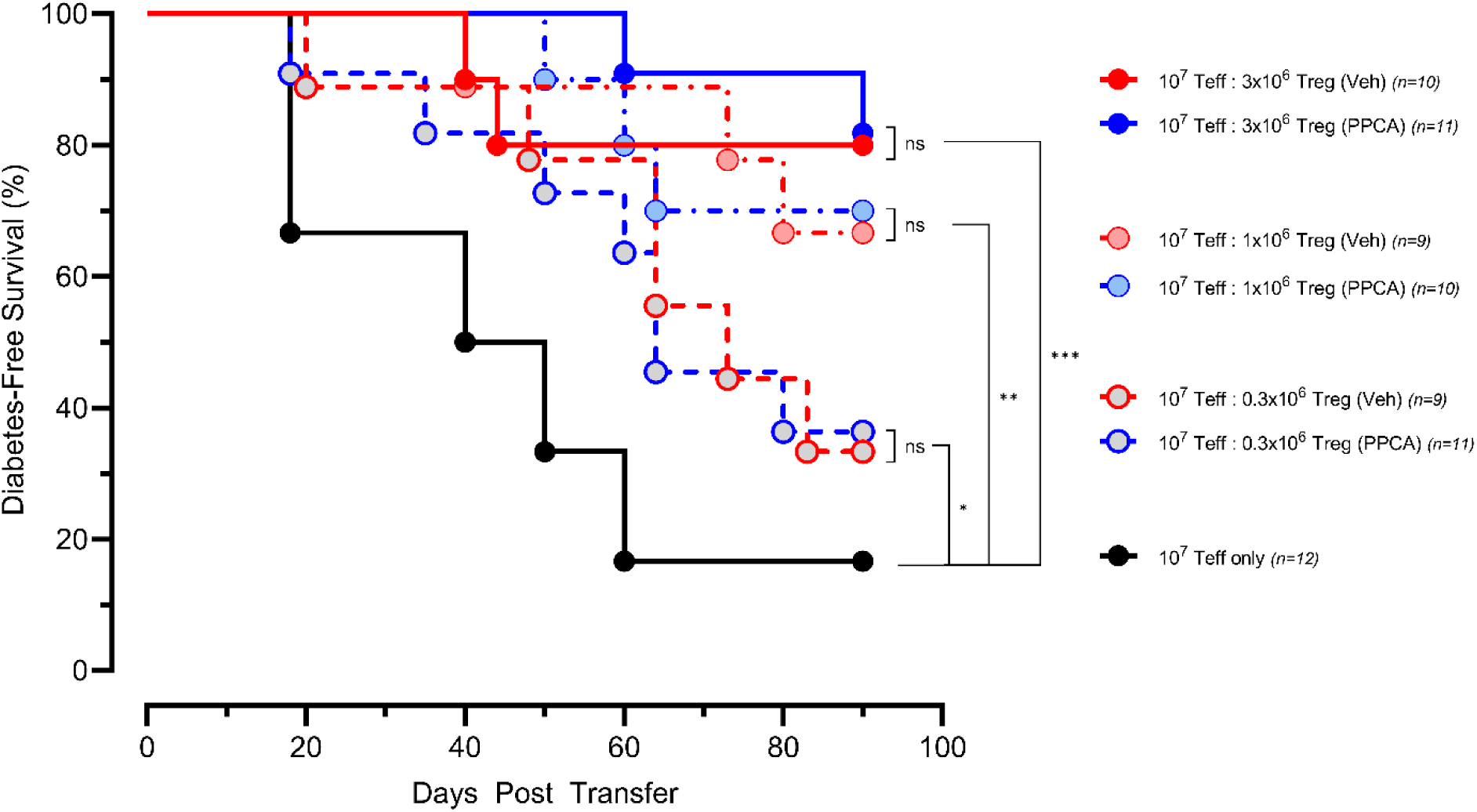
PPCA therapy does not alter the regulatory capacity of Treg on a per cell bases. Diabetic NOD.Foxp3^eGFP^ mice were treated with PPCA or vehicle and Treg cells were flow-sort purified based on eGFP expression and transferred as indicated with 10^7^ CD4^+^ NOD Teff cells into NOD.Rag1^-/-^ mice to assess regulatory capacity. No difference in regulatory capability was observed between Treg cells from either PPCA or vehicle treated mice. All three pairs, however, were significantly able to suppress the transfer of T1D when compared to the transfer of Teff cells alone. *** Denotes p<0.001; ** Denotes p<0.01; * Denotes p<0.05; ns = not significant. Disease free survival, Log-Rank (Gehan-Breslow-Wilcoxon).

PPCA therapy both CD4^+^ and CD8^+^ Teff cells, that are undergoing rapid expansion with a coincident DDR. Moreover, the more coordinated, synchronized, and robust the antigen drive, the more profound is the observed efficacy. We, therefore, reasoned that coordinated recall reactivation of memory β-cell antigen-specific CD4^+^ and CD8^+^ T cells during islet graft transplant would be an ideal moment for PPCA intervention and might result in significant long-term graft survival. To test this, we allowed a cohort of wild type, immune competent female NOD mice to develop spontaneous T1D, whereupon we treated each with subcutaneous sustained-release insulin pellets to reestablish euglycemia for ∼21 days to mimic exogenous insulin replacement therapy and to establish islet antigen-specific memory T cell subsets. As the pellet-derived insulin waned, hyperglycemia returned, and the mice were transplanted with ∼300 NOD.Rag1^-/-^ islets under the left renal capsule. This resulted in the restoration of euglycemia within 24 hours post-transplant, **Figure 5A**. We used NOD-Rag1^-/-^ donor islets as they are both autologous but lacked any passage adaptive immune lymphocytes. These islet grafts therefore mimic autologous, patient-specific iPSC-derived islets. From prior studies (24; 36), we knew that host islet-reactive lymphocytic infiltration is evident histologically by Day 3 post-transplant. Therefore, we treated the engrafted recipients with a single 3-day course of PPCA from Day 3 to Day 5 post-transplant, and thereafter monitored the mice for islet graft function by their ability to maintain non-fasting normoglycemia as assessed by standard one-step glucometer measurements, daily at first, and then weekly from 4 weeks post-transplant. As can be appreciated from **Figure 5A**, a single course of PPCA was highly efficacious in establishing long-term transplanted islet graft function in all but one of the PPCA treated recipients (cumulative graft survival, **Figure 5B**). Vehicle treated mice lost function and became hyperglycemic (mean = 18.5 days). To confirm that the islet grafts were responsible for normal glycemic control, we randomly selected 2 mice from the PPCA-treated group for graft removal by left kidney nephrectomy. In both cases the removal of the graft resulted in the return of hyperglycemia (**Sup. Figure 2**), establishing that the maintenance of euglycemia was islet graft dependent.

**Figure 5:**
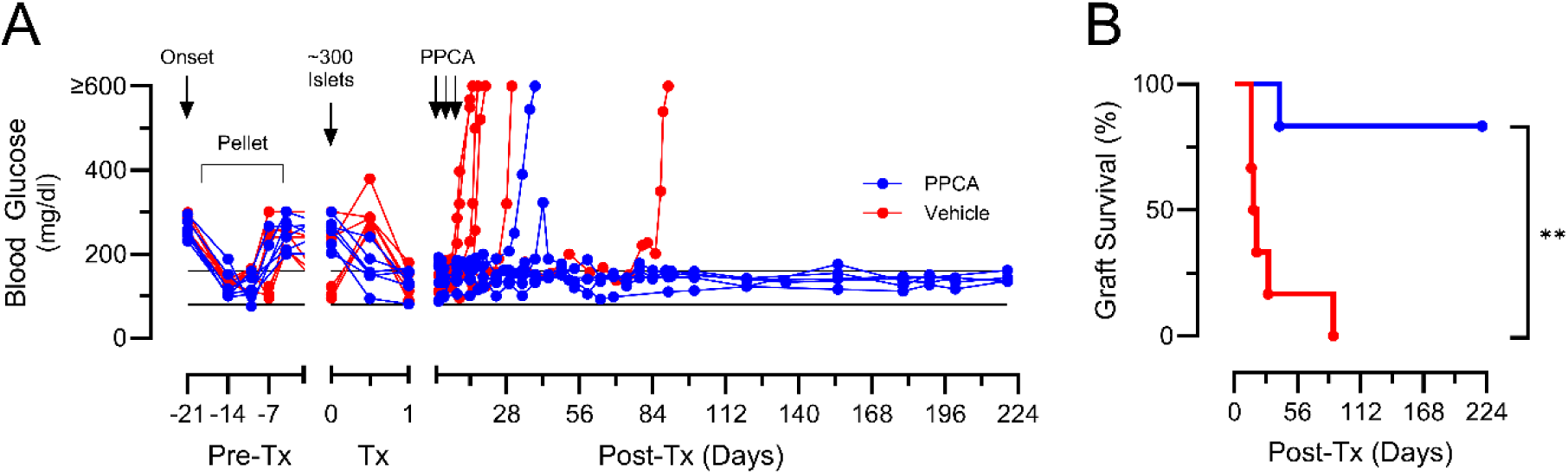
Long-term maintenance of autologous islet grafts in mice treated with a single round of PPCA therapy. **A.** NOD mice with spontaneous T1D were maintained for ∼21 days post onset using implantable insulin pellets prior to transplant of ∼300 NOD.Rag1^-/-^ islets under the left kidney capsule. On Day 3-5 post-transplant, mice were treated with either PPCA or vehicle and followed for graft survival and glycemic control. All vehicle treated mice lost graft function and glycemic control (mean = 18.5 days), n=6; while 5 out of 6 PPCA treated mice maintained glycemic control and graft function for up to 220 days post-transplant. **B.** Cumulative Graft Survival, Log-Rank, Gehan-Breslow-Wilcoxon, ** denotes p=0.0029.

To gauge the effectiveness of PPCA in purging islet-reactive T cells from islet transplanted recipients, we again transplanted diabetic NOD mice with NOD.Rag1^-/-^ islets as before and again treated them with a single course of PPCA or vehicle. After 12-14 days we removed the graft and assessed the preservation of insulin production in the graft by histology as well as determined the loss of β-cell antigen-specific T cells using tetramers to known islet antigens. As can be seen in **Figure 6**, PPCA treated mice had intact, well defined islet grafts under the kidney capsule with substantial insulin-containing β cells, as well as glucagon-containing α-cells. The vehicle treated mice, however, had markedly reduced insulin-containing β cells but did have, as expected, preservation of glucagon-containing α-cells. Flow cytometric analysis of the T cells recovered from the grafts and surrounding tissue show a substantial reduction of total CD4^+^ and CD8^+^ T cells, as well as the total numbers of CD4^+^ Teff cells, while the number of Treg cells was not statistically significantly changed, **Figure 7A**. To assess the degree of reduction in β-cell antigen-specific T cells with the graft space, we used tetramers reagents with known antigenic peptides of insulin (for CD4^+^ T cells) and IGRP (for CD8^+^ T cells). Treatment with PPCA reduced the overall numbers of CD8^+^CD44^+^ and CD4^+^CD44^+^ Teff cells (as represented in **Figure 7B**).

**Figure 6:**
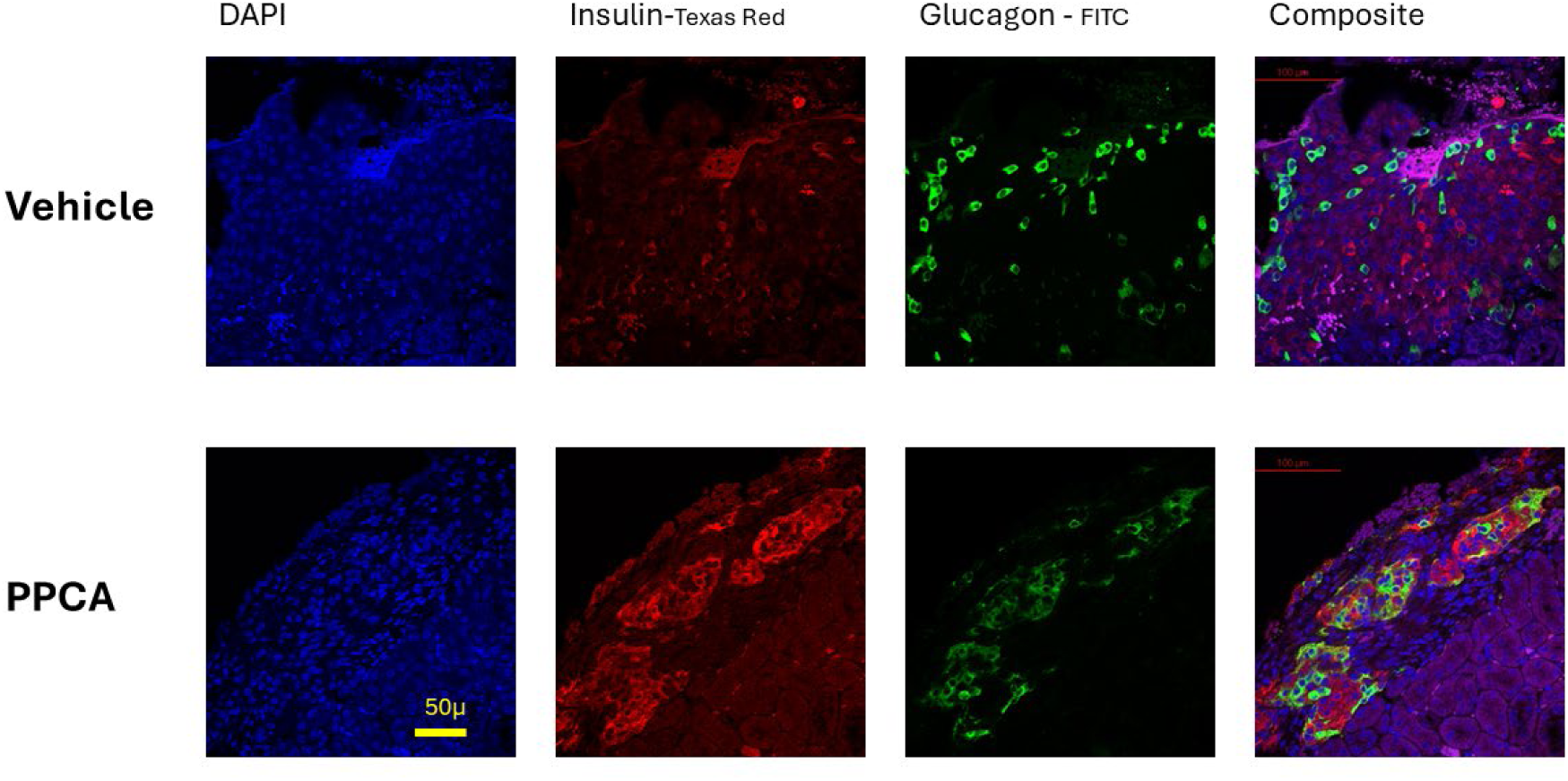
PPCA treated mice had intact, well defined islet grafts under the kidney capsule with substantial insulin-containing β cells. Engrafted mice were treated with PPCA or Vehicle as described in the text and Figure 5 on Day 3-5 post-transplant and on Day ∼12 randomly selected mice from each group were given a left nephrectomy to remove the kidney and islet graft for histology. Fluorescent microscopy shows well defined insulin producing islet grafts from PPCA treated mice. Sections were stained with anti-insulin (red) and anti-glucagon (green).

**Figure 7:**
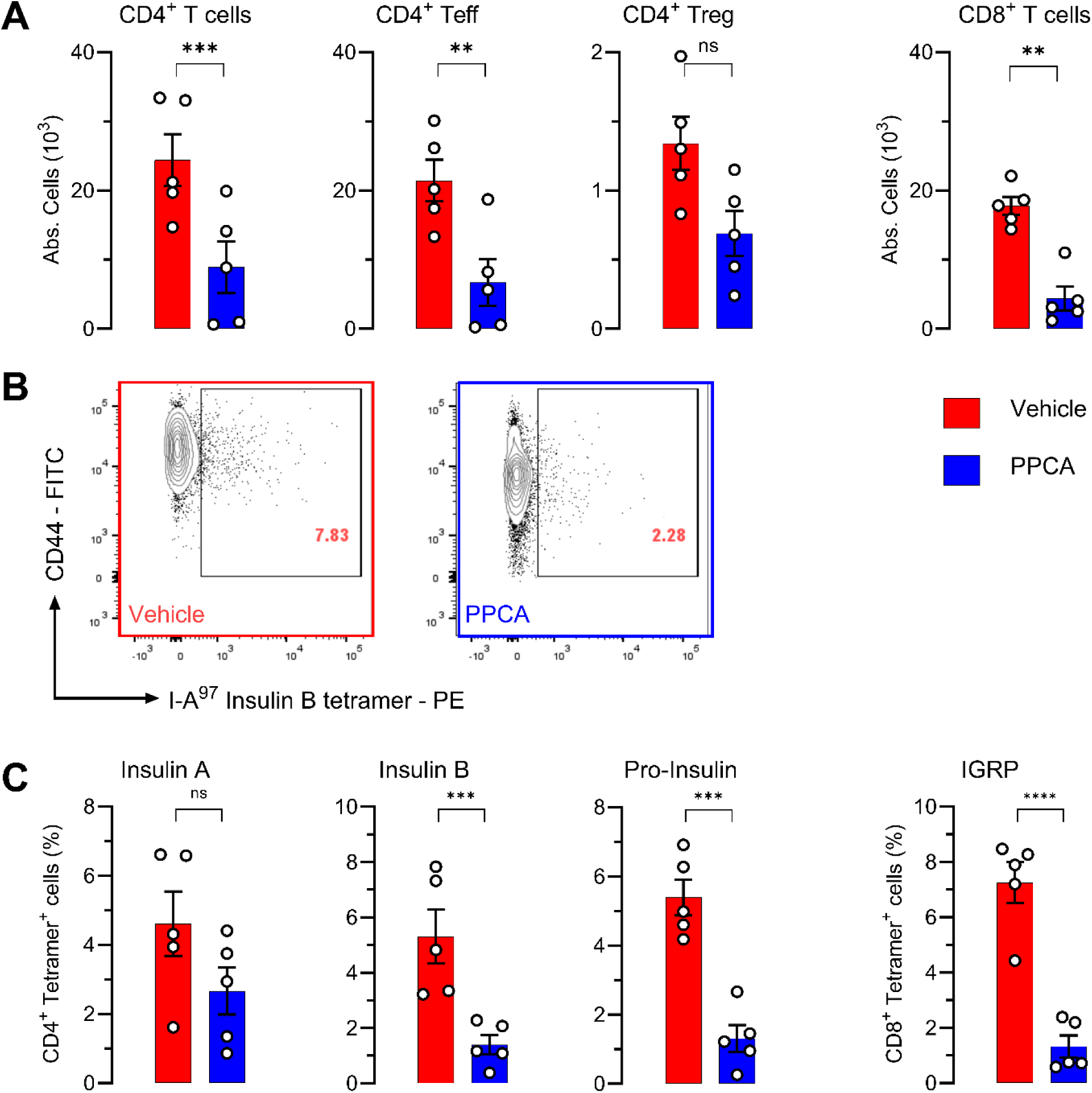
Treatment with PPCA reduced the number of graft infiltrating β-cell antigen-specific T cells. **A.** T cells from within the resected islet grafts were analyzed for the presence of total CD4^+^ and CD8^+^ as well as Teff and Treg cells. PPCA reduced overall infiltration compared to vehicle control grafts. **B.** Representative flow cytometric analysis of Insulin B chain specific CD4^+^CD44^+^ T cells in islet grafts shows reduced numbers of tetramer positive cells in PPCA treated mice. **C.** Cumulative analysis of β-cell antigen-specific T cells from PPCA and vehicle treated mice show reduced numbers of antigenic tetramer reactive T cells after PPCA treatment. ** Denotes p<0.01; *** Denotes p< 0.001; **** Denotes p<0.0001; ns= not significant. 2way ANOVA, multiple comparisons, Šídák.

Moreover, the overall number of autoantigen-specific tetramer^+^ T cells were significantly reduced (**Figure 7B** and **7C**). This was true for both I-A^g7^ class II presented antigens as well as K^d^ presented class I antigens. Thus, taken together, the overall number of CD44^+^ Teff cells as well as the measurable β cell-specific T cells were significant reduced coincident with the preservation of the islet cell grafts that could provide long-term restoration of physiological glucose regulation. Therefore, we believe that PPCA may represent an important alternative to global immunosuppression as a means of reestablishing functional tolerance to curative islet transplantation in the context of T1D.

## Discussion

Taken together, the data presented here provides a clear indication that using short-acting, small molecule inhibitors that together elevate phospho-p53 while concomitantly disrupting cell cycle control in activated autoreactive T cells leads to their apoptosis cell death that can produce long-term functional immune tolerance to autoantigens – in this case pancreatic β cell antigens – while preserving competent T regulatory function and beneficial innate and adaptive immunity (24; 25). Moreover, these data suggest that this treatment – termed PPCA therapy – is perhaps most efficacious when applied in the context of islet transplantation owing in large part to the more coordinated and robust reactivation of preexisting auto-reactive T cells thereby affording a well-defined and predictable window for treatment thereby minimizing off-target effect while preserving beneficial Treg cell function and resting T cell subsets.

Replacing β-cells via islet transplantation confers several benefits: (i) it restores natural insulin production and glucose sensing; (ii) it may prompt residual β-cells reactivation by relieving their underlying metabolic stress; and (iii) it would represent a long-term cure. Substantial advances have been made in islet transplantation and replacement (13; 14). However, the central problem of the availability of high quality islets remains, although several ongoing clinical trials (e.g., NCT02239354 and NCT03163511) are currently testing the feasibility of using iPSC-derived islets as a ready source of replacement islets (47). If produced from autologous iPSCs, one of the major hurdles – namely alloreactivity – can be avoided. However, this will still require a robust and well tolerated means of (re)-establishing functional immune tolerance to same self-antigens that drove the initial destruction of initial pancreatic β cells. This is particularly important given that existing means of exogenous insulin delivery are well established and by in large well tolerated by T1D patients. Thus, any immune suppression regime must be well tolerated with a low toxicity profile towards the islet graft and the host while providing a lower long-term risk of secondary sequalae, especially when compared to today’s existing insulin replacement modalities. Graft rejection is currently managed with lifelong immunosuppression, and although this prolongs the life of the graft it also leaves the patient susceptible to infections, cancer, and other adverse effects including some toxicity to the β-cells themselves (15; 48). While in cases of hypoglycemic unawareness, this may still be a beneficial trade-off, but for this to become a more widely available and palatable treatment, other strategies of tolerance induction are needed. PPCA therapy or similar approaches may fill this void.

As T cells transition from naïve to activated effectors to memory and then back to reactivated effectors by antigen recall they exhibit distinct features of their differentiation states, each characterized by functional and metabolic differences, distinct behavioral patterns, and varying degrees of DNA damage induction (25; 37). Full activation triggers multiple pathways in T cells, including signaling cascades activating mammalian target of rapamycin (mTOR), protein kinase C (PKC), mitogen activated protein kinases (MAPKs), and the transcription factor nuclear factor κ-light-chain-enhancer of activated B cells (NF-κB), which activate and transcribe proteins promoting cell activation, proliferation, survival, and effector functions (38). This also triggers profound metabolic changes within the cell: calcineurin signaling induces mitochondrial migration to the immunological synapse and an increase in both glycolysis and oxidative phosphorylation (38). Importantly, this process results in a cascade of increasing electron leakage due to the increased rate of oxidative phosphorylation and mitochondrial changes that leads to the formation of reactive oxygen species (ROS), which then gets transformed into hydrogen peroxide by a superoxide dismutase (MnSOD or Cu-Zn SOD) (26; 28). The ROS that are generated in this process, however, can also be harmful to the cell, and can lead to, among other things, DNA damage (25; 29; 38); thus, ROS production is the likely driver of the observed DNA damage in activated CD4^+^ and CD8^+^ Teff cells (26; 49; 50). Thus, the development of the DDR in activated T cells is inescapable and obligatory for their expansion and function. This puts p53 at the nexus of controlling the survival of these cells at the precise moment they are performing their programmed effector function. It also underpins the molecular precision by which the PPCA therapeutic approach can target and eliminate effector T cells when they are both most active and most vulnerable; this is most powerfully demonstrated when these short-acting, small molecule inhibitors are given at a moment in time when desired targets are in a state of most synchronized activation – such as shortly after (re)-exposure to self-antigen recall provided by islet transplantation.

The current technical progress towards developing a ready source of durable, functional islets for engraftment is such that it may soon represent the most likely true cure for T1D. Yet to rely on broad immunosuppression is problematic, as its drawbacks outweigh its benefits especially in those T1D patients capable of managing their disease well using existing means, as well as in the pediatric setting. It may be, then, that the best strategy is to harness and exploit processes unique to activated T effector cells; thus, paradoxically it might be more beneficial to allow or even to provoke the re-activation of deleterious autoreactive T cells to provide a clear window of opportunity to deliver therapies that drive these T cells’ apoptotic elimination. While we have not observed off-target effects of DDR targeting on quiescent central or effector memory T cell populations, our approach may be limited in T1D patients with comorbidities such as cancer, where ongoing immune responses would be beneficial. In addition, our approach may have the added benefit that abating the potency of residual Teff cells may facilitate the ability of islet specific Treg cells to reassert functional control. Further development of therapeutic strategies such as this may be the key to durable islet restoration, reversal of disease progression, and a true cure for T1D.

## Acknowledgments

The authors thank the NIH Tetramer Core for the Insulin A94-108 I-Ag7, Insulin B12-20 I-Ag7, Proinsulin47-64 I-Ag7, and IGRP206-214 H-2Kdtetramers. Additionally, we thank the CCHMC Pathology Core for assistance in processing histological samples, the University of Cincinnati Genomics, Epigenomics, and Sequencing Core for assistance with RNA sequencing.

Single Cell RNA-Seq datasets have been deposited Datasets have been with the following accession number: GSE262644 at: https://www.ncbi.nlm.nih.gov/geo/query/acc.cgi?acc=GSE262644

## Author Contributions

E.E.C., D.H.P., J.D.K. and M.K. performed experiments and acquired data. M.K., A.B., and J.D.K. interpreted and analyzed data. M.B.J., and J.D.K. contributed to conceptual and research design. J.D.K. wrote the manuscript. J.D.K. is the guarantor of this work and, as such, had full access to all the data in the study and takes responsibility for the integrity of the data and the accuracy of the data analysis.

## References

1. DiMeglio LA, Evans-Molina C, Oram RA: Type 1 diabetes. Lancet 2018;391:2449–2462

2. Miller KM, Foster NC, Beck RW, Bergenstal RM, DuBose SN, DiMeglio LA, Maahs DM, Tamborlane WV, Network TDEC: Current state of type 1 diabetes treatment in the U.S.: updated data from the T1D Exchange clinic registry. Diabetes Care 2015;38:971–978

3. Andre I, Gonzalez A, Wang B, Katz J, Benoist C, Mathis D: Checkpoints in the progression of autoimmune disease: lessons from diabetes models. Proc Natl Acad Sci U S A 1996;93:2260–2263

4. Katz JD, Benoist C, Mathis D: T helper cell subsets in insulin-dependent diabetes. Science 1995;268:1185–1188

5. Wang B, Andre I, Gonzalez A, Katz JD, Aguet M, Benoist C, Mathis D: Interferon-gamma impacts at multiple points during the progression of autoimmune diabetes. Proc Natl Acad Sci U S A 1997;94:13844–13849

6. Schneider A, Rieck M, Sanda S, Pihoker C, Greenbaum C, Buckner JH: The effector T cells of diabetic subjects are resistant to regulation via CD4+ FOXP3+ regulatory T cells. J Immunol 2008;181:7350–7355

7. McClymont SA, Putnam AL, Lee MR, Esensten JH, Liu W, Hulme MA, Hoffmuller U, Baron U, Olek S, Bluestone JA, Brusko TM: Plasticity of human regulatory T cells in healthy subjects and patients with type 1 diabetes. J Immunol 2011;186:3918–3926

8. Long SA, Buckner JH: CD4+FOXP3+ T regulatory cells in human autoimmunity: more than a numbers game. J Immunol 2011;187:2061–2066

9. Buckner JH: Mechanisms of impaired regulation by CD4(+)CD25(+)FOXP3(+) regulatory T cells in human autoimmune diseases. Nat Rev Immunol 2010;10:849–859

10. Ehlers MR: Immune interventions to preserve beta cell function in type 1 diabetes. J Investig Med 2016;64:7–13

11. Attias M, Al-Aubodah T, Piccirillo CA: Mechanisms of human FoxP3(+) T(reg) cell development and function in health and disease. Clin Exp Immunol 2019;197:36–51

12. Krogvold L, Skog O, Sundstrom G, Edwin B, Buanes T, Hanssen KF, Ludvigsson J, Grabherr M, Korsgren O, Dahl-Jorgensen K: Function of Isolated Pancreatic Islets From Patients at Onset of Type 1 Diabetes: Insulin Secretion Can Be Restored After Some Days in a Nondiabetogenic Environment In Vitro: Results From the DiViD Study. Diabetes 2015;64:2506–2512

13. Cito M, Pellegrini S, Piemonti L, Sordi V: The potential and challenges of alternative sources of beta cells for the cure of type 1 diabetes. Endocr Connect 2018;7:R114–R125

14. Aguayo-Mazzucato C, Bonner-Weir S: Pancreatic beta Cell Regeneration as a Possible Therapy for Diabetes. Cell Metab 2018;27:57–67

15. Barton FB, Rickels MR, Alejandro R, Hering BJ, Wease S, Naziruddin B, Oberholzer J, Odorico JS, Garfinkel MR, Levy M, Pattou F, Berney T, Secchi A, Messinger S, Senior PA, Maffi P, Posselt A, Stock PG, Kaufman DB, Luo X, Kandeel F, Cagliero E, Turgeon NA, Witkowski P, Naji A, O’Connell PJ, Greenbaum C, Kudva YC, Brayman KL, Aull MJ, Larsen C, Kay TW, Fernandez LA, Vantyghem MC, Bellin M, Shapiro AM: Improvement in outcomes of clinical islet transplantation: 1999-2010. Diabetes Care 2012;35:1436–1445

16. Maxwell KG, Millman JR: Applications of iPSC-derived beta cells from patients with diabetes. Cell Rep Med 2021;2:100238

17. Du Y, Liang Z, Wang S, Sun D, Wang X, Liew SY, Lu S, Wu S, Jiang Y, Wang Y, Zhang B, Yu W, Lu Z, Pu Y, Zhang Y, Long H, Xiao S, Liang R, Zhang Z, Guan J, Wang J, Ren H, Wei Y, Zhao J, Sun S, Liu T, Meng G, Wang L, Gu J, Wang T, Liu Y, Li C, Tang C, Shen Z, Peng X, Deng H: Human pluripotent stem-cell-derived islets ameliorate diabetes in non-human primates. Nat Med 2022;28:272–282

18. Warshauer JT, Bluestone JA, Anderson MS: New Frontiers in the Treatment of Type 1 Diabetes. Cell Metab 2020;31:46–61

19. Malmegrim KC, de Azevedo JT, Arruda LC, Abreu JR, Couri CE, de Oliveira GL, Palma PV, Scortegagna GT, Stracieri AB, Moraes DA, Dias JB, Pieroni F, Cunha R, Guilherme L, Santos NM, Foss MC, Covas DT, Burt RK, Simoes BP, Voltarelli JC, Roep BO, Oliveira MC: Immunological Balance Is Associated with Clinical Outcome after Autologous Hematopoietic Stem Cell Transplantation in Type 1 Diabetes. Front Immunol 2017;8:167

20. Marino E, Richards JL, McLeod KH, Stanley D, Yap YA, Knight J, McKenzie C, Kranich J, Oliveira AC, Rossello FJ, Krishnamurthy B, Nefzger CM, Macia L, Thorburn A, Baxter AG, Morahan G, Wong LH, Polo JM, Moore RJ, Lockett TJ, Clarke JM, Topping DL, Harrison LC, Mackay CR: Gut microbial metabolites limit the frequency of autoimmune T cells and protect against type 1 diabetes. Nat Immunol 2017;18:552–562

21. Bednar KJ, Tsukamoto H, Kachapati K, Ohta S, Wu Y, Katz JD, Ascherman DP, Ridgway WM: Reversal of New-Onset Type 1 Diabetes With an Agonistic TLR4/MD-2 Monoclonal Antibody. Diabetes 2015;64:3614–3626

22. Foda BM, Ciecko AE, Serreze DV, Ridgway WM, Geurts AM, Chen YG: The CD137 Ligand Is Important for Type 1 Diabetes Development but Dispensable for the Homeostasis of Disease-Suppressive CD137(+) FOXP3(+) Regulatory CD4 T Cells. J Immunol 2020;204:2887–2899

23. Itoh A, Ortiz L, Kachapati K, Wu Y, Adams D, Bednar K, Mukherjee S, Chougnet C, Mittler RS, Chen YG, Dolan L, Ridgway WM: Soluble CD137 Ameliorates Acute Type 1 Diabetes by Inducing T Cell Anergy. Front Immunol 2019;10:2566

24. Carroll KR, Elfers EE, Stevens JJ, McNally JP, Hildeman DA, Jordan MB, Katz JD: Extending Remission and Reversing New-Onset Type 1 Diabetes by Targeted Ablation of Autoreactive T Cells. Diabetes 2018;67:2319–2328

25. McNally JP, Millen SH, Chaturvedi V, Lakes N, Terrell CE, Elfers EE, Carroll KR, Hogan SP, Andreassen PR, Kanter J, Allen CE, Henry MM, Greenberg JN, Ladisch S, Hermiston ML, Joyce M, Hildeman DA, Katz JD, Jordan MB: Manipulating DNA damage-response signaling for the treatment of immune-mediated diseases. Proc Natl Acad Sci U S A 2017;114:E4782–E4791

26. Li KP, Shanmuganad S, Carroll K, Katz JD, Jordan MB, Hildeman DA: Dying to protect: cell death and the control of T-cell homeostasis. Immunol Rev 2017;277:21–43

27. Elbaek CR, Petrosius V, Sorensen CS: WEE1 kinase limits CDK activities to safeguard DNA replication and mitotic entry. Mutat Res 2020;819-820:111694

28. Carroll KR, Katz JD: Restoring tolerance to beta-cells in Type 1 diabetes: Current and emerging strategies. Cell Immunol 2022;380:104593

29. McNally JP, Elfers EE, Terrell CE, Grunblatt E, Hildeman DA, Jordan MB, Katz JD: Eliminating encephalitogenic T cells without undermining protective immunity. J Immunol 2014;192:73–83

30. Zheng GX, Terry JM, Belgrader P, Ryvkin P, Bent ZW, Wilson R, Ziraldo SB, Wheeler TD, McDermott GP, Zhu J, Gregory MT, Shuga J, Montesclaros L, Underwood JG, Masquelier DA, Nishimura SY, Schnall-Levin M, Wyatt PW, Hindson CM, Bharadwaj R, Wong A, Ness KD, Beppu LW, Deeg HJ, McFarland C, Loeb KR, Valente WJ, Ericson NG, Stevens EA, Radich JP, Mikkelsen TS, Hindson BJ, Bielas JH: Massively parallel digital transcriptional profiling of single cells. Nat Commun 2017;8:14049

31. Kotliar M, Kartashov A, Barski A: Accelerating Single-Cell Sequencing Data Analysis with SciDAP: A User-Friendly Approach. bioRxiv 2024;

32. Stuart T, Butler A, Hoffman P, Hafemeister C, Papalexi E, Mauck WM, 3rd, Hao Y, Stoeckius M, Smibert P, Satija R: Comprehensive Integration of Single-Cell Data. Cell 2019;177:1888–1902 e1821

33. Subramanian A, Tamayo P, Mootha VS, Mesirov JP: Gene set enrichment analysis: A knowledge-based approach for interpreting genome-wide expression profiles. Proc Natl Acad Sci U S A 2005;102:15545–15550

34. Liberzon A, Birger C, Thorvaldsdottir H, Ghandi M, Mesirov JP, Tamayo P: The Molecular Signatures Database (MSigDB) hallmark gene set collection. Cell Syst 2015;1:417–425

35. Durinck S, Spellman PT, Birney E, Huber W: Mapping identifiers for the integration of genomic datasets with the R/Bioconductor package biomaRt. Nat Protoc 2009;4:1184–1191

36. Pakala SV, Chivetta M, Kelly CB, Katz JD: In autoimmune diabetes the transition from benign to pernicious insulitis requires an islet cell response to tumor necrosis factor alpha. J Exp Med 1999;189:1053–1062

37. Buck MD, O’Sullivan D, Klein Geltink RI, Curtis JD, Chang CH, Sanin DE, Qiu J, Kretz O, Braas D, van der Windt GJ, Chen Q, Huang SC, O’Neill CM, Edelson BT, Pearce EJ, Sesaki H, Huber TB, Rambold AS, Pearce EL: Mitochondrial Dynamics Controls T Cell Fate through Metabolic Programming. Cell 2016;166:63–76

38. Park BV, Pan F: Metabolic regulation of T cell differentiation and function. Mol Immunol 2015;68:497–506

39. Pakala SV, Kurrer MO, Katz JD: T helper 2 (Th2) T cells induce acute pancreatitis and diabetes in immune-compromised nonobese diabetic (NOD) mice. J Exp Med 1997;186:299–306

40. Liu X, Li H, Zhong B, Blonska M, Gorjestani S, Yan M, Tian Q, Zhang DE, Lin X, Dong C: USP18 inhibits NF-kappaB and NFAT activation during Th17 differentiation by deubiquitinating the TAK1-TAB1 complex. J Exp Med 2013;210:1575–1590

41. Yang L, Jing Y, Kang D, Jiang P, Li N, Zhou X, Chen Y, Westerberg LS, Liu C: Ubiquitin-specific peptidase 18 regulates the differentiation and function of Treg cells. Genes Dis 2021;8:344–352

42. Fensterl V, Sen GC: The ISG56/IFIT1 gene family. J Interferon Cytokine Res 2011;31:71–78

43. Xin Y, He Z, Mei Y, Li X, Zhao Z, Zhao M, Yang M, Wu H: Interferon-alpha regulates abnormally increased expression of RSAD2 in Th17 and Tfh cells in systemic lupus erythematosus patients. Eur J Immunol 2023;53:e2350420

44. Chwetzoff S, d’Andrea S: Ubiquitin is physiologically induced by interferons in luminal epithelium of porcine uterine endometrium in early pregnancy: global RT-PCR cDNA in place of RNA for differential display screening. FEBS Lett 1997;405:148–152

45. Lee Y, Awasthi A, Yosef N, Quintana FJ, Xiao S, Peters A, Wu C, Kleinewietfeld M, Kunder S, Hafler DA, Sobel RA, Regev A, Kuchroo VK: Induction and molecular signature of pathogenic TH17 cells. Nat Immunol 2012;13:991–999

46. Lee YK, Turner H, Maynard CL, Oliver JR, Chen D, Elson CO, Weaver CT: Late developmental plasticity in the T helper 17 lineage. Immunity 2009;30:92–107

47. Zhao K, Shi Y, Yu J, Yu L, Kohler M, Mael A, Kolton A, Joyce T, Odorico J, Berggren PO, Yang SN: In Vivo Ca(V)3 Channel Inhibition Promotes Maturation of Glucose-Dependent Ca(2+) Signaling in Human iPSC-Islets. Biomedicines 2023;11

48. Berney T, Andres A, Toso C, Majno P, Squifflet JP: mTOR Inhibition and Clinical Transplantation: Pancreas and Islet. Transplantation 2018;102:S30–S31

49. Sena LA, Li S, Jairaman A, Prakriya M, Ezponda T, Hildeman DA, Wang CR, Schumacker PT, Licht JD, Perlman H, Bryce PJ, Chandel NS: Mitochondria are required for antigen-specific T cell activation through reactive oxygen species signaling. Immunity 2013;38:225–236

50. Cameron AM, Castoldi A, Sanin DE, Flachsmann LJ, Field CS, Puleston DJ, Kyle RL, Patterson AE, Hassler F, Buescher JM, Kelly B, Pearce EL, Pearce EJ: Inflammatory macrophage dependence on NAD(+) salvage is a consequence of reactive oxygen species-mediated DNA damage. Nat Immunol 2019;20:420–432

