## Supplemental Files for "Long-term Tolerance to Islet Transplantation via Targeted Reduction of beta cell-specific T cells"

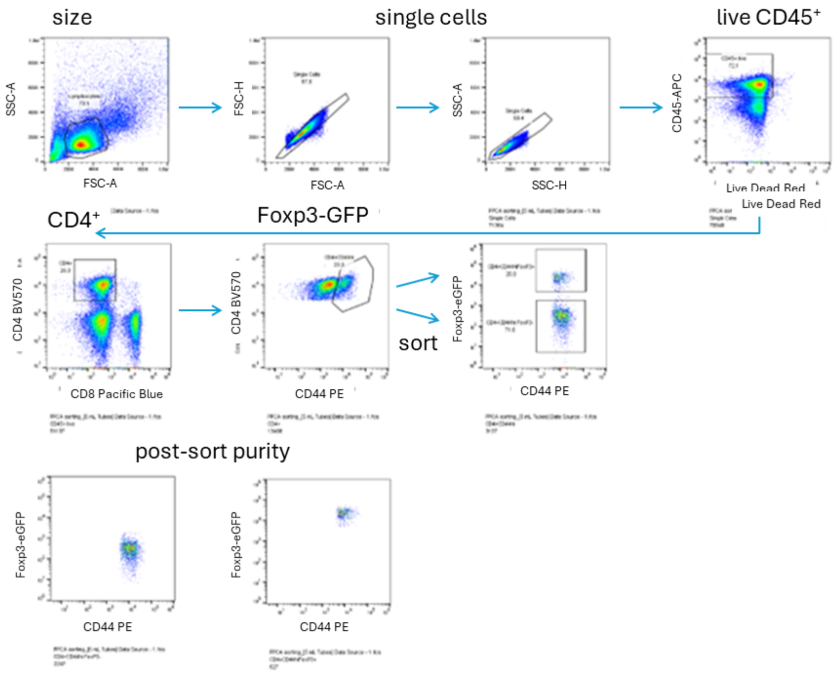


**Sup. Figure 1: Sort Purification strategy for Teff and Treg cells from NOD.Foxp3$^{eGFP}$ mice.** Diabetic NOD.Foxp3$^{eGFP}$ mice treated with vehicle or PPCA for 3 consecutive days and then the splenic and pancreatic lymph node cells were harvested, and sort purified as shown on a Sony MA900 flow sorter. Lymphocytes were gated on size and granularity via SSC-A vs FSC-A, then on singlets via FSC-A vs FSC-H followed by CD45$^{+}$. CD4$^{+}$ cells were identified via CD4 vs CD8 of CD45$^{+}$ gated cells. CD44$^{hi}$CD4$^{+}$ from the CD4+ gated population were then gated into Treg (CD44$^{hi}$Fop3$^{eGFP+}$) and Teff (CD44$^{hi}$Foxp3$^{eGFP-}$) and sorted into individual tubes. Sort purity was confirmed by reanalysis of sorted populations. Sort purified cells were then enumerated, mixed as described in the text and Figure 4 and injected into NOD.Rag1$^{-/-}$ mice.

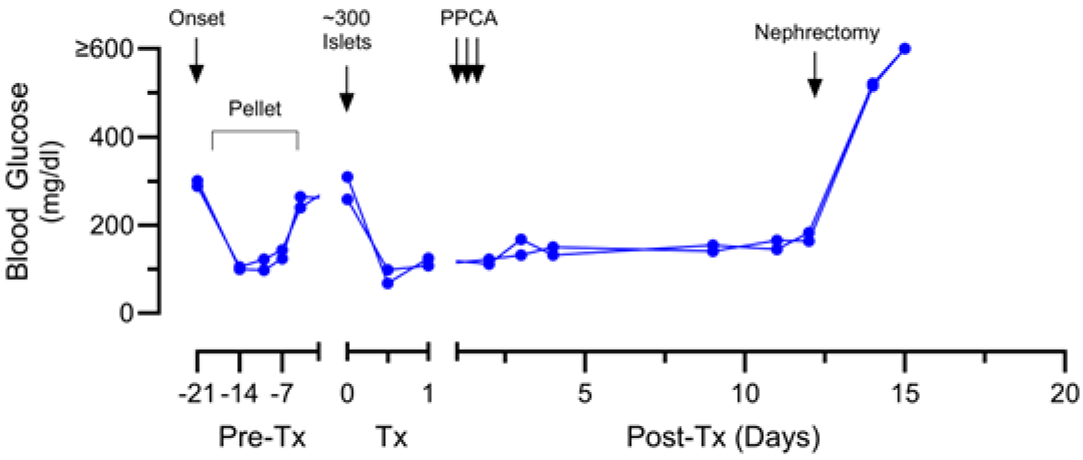


**Sup. Figure 2: Resection of engrafted islets via a left nephrectomy results in loss of euglycemia and demonstrates that the islet grafts were functional and provided normal blood glucose regulation.** NOD mice with spontaneous T1D were maintained for ~21 days post onset using implantable insulin pellets prior to transplant of ~300 NOD.Rag1$^{-/-}$ islets under the left kidney capsule. On Day 3-5 post-transplant, mice were treated with PPCA and followed for graft survival and glycemic control.  On day 12 post-transplant, 2 mice were randomly selected to undergo the nephrectomy of the left (engrafted) kidney
